# Codon usage bias in animals: disentangling the effects of natural selection, effective population size and GC-biased gene conversion

**DOI:** 10.1101/184283

**Authors:** N. Galtier, C. Roux, M. Rousselle, J. Romiguier, E. Figuet, S. Glémin, N. Bierne, L. Duret

## Abstract

Selection on codon usage bias is well documented in a number of microorganisms. Whether codon usage is also generally shaped by natural selection in large organisms, despite their relatively small effective population size (*N_e_*), is unclear. Codon usage bias in animals has only been studied in a handful of model organisms so far, and can be affected by confounding, non-adaptive processes such as GC-biased gene conversion and experimental artefacts. Using population transcriptomics data we analysed the relationship between codon usage, gene expression, allele frequency distribution and recombination rate in 31 non-model species of animals, each from a different family, covering a wide range of effective population sizes. We disentangled the effects of translational selection and GC-biased gene conversion on codon usage by separately analysing GC-conservative and GC-changing mutations. We report evidence for effective translational selection on codon usage in large-*N_e_* species of animals, but not in small-*N_e_* ones, in agreement with the nearly neutral theory of molecular evolution. C- and T-ending codons are generally preferred over synonymous G- and A-ending ones, for reasons that remain to be determined. In contrast, we uncovered a conspicuous effect of GC-biased gene conversion, which is widespread in animals and the main force determining the fate of AT↔GC mutations. Intriguingly, the strength of its effect was uncorrelated with *N_e_*.

## Background

The reasons why synonymous codons do not occur at equal frequencies in protein coding sequences have been puzzling molecular evolutionary researchers for decades (Duret 2002, Hershberg and Petrov 2008). One fascinating aspect is the early discovery that, in various microbial genomes, codon usage responds to natural selection. In *Escherichia coli* and *Saccharomyces cerevisiae*, for instance, the codons most commonly observed in highly expressed genes match the most abundant tRNAs in the cell (Ikemura 1985), strongly suggesting that codon usage and tRNA content have co-evolved in a way that optimizes translation – hence the term “translational selection”. These observations promoted codon usage bias as a textbook example of a weak selective pressure operating at molecular level, detectable from patterns of coding sequence variation, but difficult to apprehend experimentally.

Because selection on codon usage is presumably weak, other evolutionary forces might contribute to explain its variation across genes and genomes (Sharp and Li 1986). In particular, it is expected that, in small populations, the random fluctuations of allele frequencies due to genetic drift should decrease the efficiency of natural selection and lessen the efficiency of selection on codon usage. The theory predicts that if *N_e_*, the effective population size, is sufficiently small such that the 4*N_e_s* product is much lower than 1, *s* being the selection coefficient in favour of optimal codons, then the effect of selection should be negligible. One question of interest, therefore, is whether selection shapes codon usage in large organisms, such as animals, the same way as in microbes, despite their presumably smaller *N_e_*.

Evidence for selection on codon usage has been reported in fruit flies (Shields et al. 1988, Akashi 1994, Bierne & Eyre-Walker 2006), in the nematode *Caenorhabditis elegans* (Duret and Mouchiroud 1999), and in the branchiopod *Daphnia pulex* (Lynch et al. 2017). In contrast, codon usage in mammals is primarily governed by within-genome variation in GC-content, and only weakly, if any, correlated to gene expression and tRNA content (Sémon et al. 2006, Rudolph et al. 2016, Pouyet et al. 2017). The effectiveness of selection on codon usage in small-sized invertebrates but not in large-sized vertebrates is superficially in agreement with the hypothesis of a *N_e_* effect. Please note, however, that the evidence so far relies on a relatively small number of species of animals.

Subramanian (2008) analysed codon usage bias intensity across 20 species of eukaryotes and reported a higher bias in short generation time, presumably large-*N_e_* species than in long generation time, presumably small-*N_e_* ones. This author questioned the *N_e_* hypothesis and rather suggested that *s* might vary among species. According to his hypothesis, the selective pressure on translation efficiency would be stronger in fast growing species. Interestingly, growth rate seems to be the main determinant of among-species variation in codon usage bias intensity in bacteria (Sharp et al. 2005, Rocha 2004, Vieira-Silva et al 2011). On the other hand, Machado et al. (2017) argued that *s* is not necessarily constant across synonymous codons and mutations. Comparison of polymorphism and divergence patterns in *Drosophila melanogaster* indeed suggested that both strong (4*N_e_s*>>1) and weak (4*N_e_s*~1) selection applies on synonymous sites in this species (Lawrie et al. 2013, Machado et al. 2017). Is is also well established that in *E. coli* and *S. cerevisiae* selection for optimal translation is stronger in highly expressed that in lowly expressed genes, and even in humans there is documented evidence for a phenotypic effect of specific synonymous mutations (Sauna & Kimchi-Sarfaty 2011). The question “does codon usage affect translational efficiency in species X” should therefore probably be rephrased as “what fraction of synonymous mutations in species X is effectively selected?”, and “how does *N_e_* influence this fraction?”

Besides selection and drift, patterns of codon usage might be influenced by neutral, directional forces such as mutation biases and GC-biased gene conversion (gBGC, Duret & Galtier 2009, Mugal et al. 2015). gBGC is a recombination-associated segregation bias that favours G and C over A and T alleles in high recombining regions. The existence of gBGC has been experimentally demonstrated in yeast (Mancera et al. 2008, Lesecque et al. 2013), humans (Williams et al. 2015, Halldorsson et al. 2016), flycatcher (Smeds et al. 2016) and *Daphnia* (Keith et al. 2016). gBGC has been identified as the main driver of GC-content evolution in vertebrates (Duret & Galtier 2009, Figuet et al. 2014, Bolivar et al. 2016, Glémin et al. 2015) and several other taxa (Pessia et al. 2012, Glémin et al. 2014, Wallberg et al. 2015). In many respects the expected impact of gBGC on patterns of sequence variation is similar to that of directional selection. For instance, the expected fate and frequency distribution of an allele promoted by gBGC is identical to that of a favourable allele under codominant selection (Nagylaky 1983). Importantly, the so-called “preferred” synonymous codons – i.e., codons more frequently used in high-expressed genes – often end with C or G, *D. melanogaster* being an extreme example in which all the preferred codons are C- or G-ending (Duret & Mouchiroud 1999). This implies that the effects of translational selection and gBGC on synonymous positions can be very difficult to disentangle from coding sequence analysis only (Jackson et al. 2017, but see Clément et al. 2017). gBGC, however, is expected to apply more strongly to high recombining regions, and to affect non coding, flanking sequences as well as coding ones, unlike translational selection.

Preferred codons are usually defined as codons used more frequently in high-expressed than in low-expressed genes (e.g. Duret and Mouchiroud 1999). This could be problematic in practice because DNA or cDNA libraries generated for high throughput sequencing are known to be biased with respect to sequence base composition (Dohm et al. 2008, Aird et al. 2011). The GC-richest and GC-poorest fractions of target DNA are typically under-represented, and sequences of medium GC-content over-represented, in Illumina data (Choudhari & Grigoriev 2017). This experimental bias, if not properly taken into account, could corrupt the definition of preferred codons in generating false correlations between codon usage and sequencing coverage. The bias apparently varies between experiments and libraries, which makes it difficult to model and correct for (Benjamini & Speed 2012). GC-content is therefore a potential confounder of analyses of selection on codon usage, both biologically and methodologically.

Current knowledge on codon usage biases in animals is therefore limited in at least two respects. First, published analyses have so far focused on a relatively small number of species – mainly model organisms – and an even smaller number of taxa – mainly drosophilids and vertebrates. Secondly, the confounding effects of gBGC and experimental biases have not always been taken into account. We therefore lack a global picture of the relative impact of selection, drift and gBGC on codon usage evolution in animals. Here we analysed a data set covering 31 non-model species of animals. In each species the transcriptome of five to eleven diploid individuals plus one outgroup have been previously sequenced (Romiguier et al. 2014). In principle transcriptome-based population genomic data are ideal for codon usage bias analysis in providing access to codon usage tables, gene expression level, allele frequencies at polymorphic positions, and flanking UTR sequences. We focused our analysis of translational selection on GC-conservative pairs of synonymous codons – i.e., codons differing by a G↔C or an A↔T substitution – and separately analysed the effect of gBGC. We found that translational selection on codon usage is only detectable in short-lived, large-*N_e_* species, whereas gBGC is widespread across animals and of strength apparently independent of *N_e_*.

## Methods

### Species sampling

We used recently published Illumina transcriptome data from population samples of non model animals (Romiguier et al. 2014, Rousselle et al. 2016, Ballenghien et al. 2017, Romiguier et al. 2017), which covered 32 distinct families of Metazoa. In each family we selected the species with the largest number of individuals, provided this number was five or more. Mosquito *Culex pipiens* (Culicidae) was excluded because transcriptome assembly in this species yielded a small number of very short contigs (Romiguier et al. 2014). Harvester ant *Messor barbarus* (Formicidae) was excluded because of its peculiar mating system, which dramatically departs the Hardy-Weinberg assumption (Romiguier et al. 2017). Its sister species *Messor capitatus* was rather included, despite a lower number of sampled individuals. The final data set included 31 species, of which seven vertebrates, six insects, five molluscs, three crustaceans, three echinoderms, two tunicates, two annelids, one nematode, one nemertian and one cnidarian (Table S1). Five to eleven individuals per species were analysed. For each of these focal species one outgroup from the same family was selected. The tissues from which RNA has been extracted, which differ across species, are provided in Table S1.

Romiguier et al. (2014) reported significant correlations between life history traits, such as species longevity, fecundity and propagule size, and population genomic variables theoretically related to *N_e_*, such as the synonymous diversity, π_S_, and the ratio of nonsynonymous over synonymous heterozygosity, π_N_/π_S_. In this study we used longevity, propagule size and π_N_/π_S_ as markers of the long-term *N_e_* of the analysed species. π_N_/π_S_ is expected to be negatively correlated with *N_e_* due to the decreased efficiency of purifying selection against slightly deleterious non-synonymous alleles in small populations (Lanfear et al. 2014).

### Transcriptome assembly and annotation

Transcriptome assembly, open reading frame (ORF) prediction, orthology prediction and alignment between focal and outgroup coding sequences were achieved using the Abyss, Cap3, Trinity_ORF, BLAST and MACSe programs, as previously described (Gayral et al. 2013, Romiguier et al. 2014). We only retained contigs containing a predicted coding sequences (CDS) longer than 200 bp. The median number of contigs per species was 3480 (Table S1). Contig expression level was measured as the per base pair read depth. For each contig of each species, we calculated GC-content at first and second codon positions (GC12), third codon positions (GC3) and UTR (GC_UTR), and the frequency of the 61 sense codons.

### SNP and genotype calling

Genotypes and single nucleotide polymorphisms (SNPs) were called using the reads2snp program, which was designed for genotyping based on RNAseq data (Tsagkogeorga et al. 2012, Gayral et al. 2013, Ballenghien et al. 2017). This method models read counts at each position as a multinomial distribution determined by allele frequencies, genotype frequencies, sequencing error rate and cross-contamination rate. Allele frequencies are estimated a priori from read counts across all individuals. Genotype frequencies are assumed to follow the Hardy-Weinberg prior. The method first estimates the error rate by maximum likelihood, then the posterior distribution of genotypes in the empirical Bayesian framework (Tsagkogeorga et al. 2012). Contamination rate (Flickinger et al. 2015, Ballenghien et al. 2017) was here set to 0.2. This parameter likely captures a combination of effects leading to overdispersion of read counts and spurious calls of heterozygote genotypes (Ballenghien et al. 2017). A filter for false SNPs due to hidden paralogy was applied posterior to genotyping.

### Site frequency spectrum analysis

Synonymous SNPs were oriented assuming that the state observed at the orthologous position in the outgroup is ancestral. Unfolded site frequency spectra (SFS), i.e., the observed distribution of derived allele frequency across SNPs, were built separately for AT→GC, GC→AT, and A↔T or G↔C synonymous mutations. The three SFS’s were analysed in the maximum likelihood framework using model M1 in Glémin et al. (2015), which accounts for the effect of mutation bias, gBGC, drift and SNP orientation error. The significance of gBGC was assessed by comparing the full model to a model assuming no gBGC via a likelihood ratio test.

### Effect of recombination rate

We approached the population-scaled recombination rate of each locus via a calculation based on the four-gamete rule (Hudson 1985). For every pair of SNPs in a locus, haplotypes were identified from individuals homozygous at both SNPs, and from individuals heterozygous at one SNP and homozygous at the other SNP. In these two situations, linkage relationship between alleles can be determined with certainty even from unphased data. Individuals carrying a heterozygous genotype at both SNPs were here disregarded. When the four possible haplotypes were found to be segregating in the sample, a recombination event was inferred. The total number of recombination events per contig was recorded by summing across pairs of SNPs, taking care of only counting once events supported by non independent pairs of SNPs. We defined *R*_fgr_ (for “four-gamete rule”) as the ratio of total number of inferred recombination events by contig length. This was calculated, in each species, for each contig carrying at least one pair of SNPs, excluding singletons, such that four haplotypes or more could be inferred. Contigs departing these conditions were not considered eligible for recombination analysis, and missing data was recorded. Species in which less than 500 eligible contigs were available were not considered.

### Insect transcriptome data

We downloaded from the NCBI SRA database Illumina RNAseq reads from 50 species of insects – 20 eusocial and 30 solitary. Specifically, we selected five species of ant, five eusocial bees, five eusocial wasps, five termites, five solitary hymenopterans, five solitary cockroaches (Blattodea, same order as termites), and 20 species from other orders of insects (Table S2). Species sampled in the context of the 1KITE project (http://www.1kite.org/) were favoured whenever possible. Transcriptome assemblies were downloaded from NCBI Sequence Set in 43 species. In the remaining seven species transcriptomes were assembled as above. In each species reads were mapped to predicted cDNA. Depth of coverage and codon usage were computed for each contig of each species. The species list and accession numbers are provided in Table S2.

## Results

### GC-changing vs. GC-conservative mutations

In each of the 31 focal species we correlated GC12, GC3 and GC_UTR across genes. The three measures of GC-content were strongly correlated with each other in nearly all species (Table S1). We calculated the mean allele frequency of the G or C allele at synonymous, AT↔GC SNPs. The mean frequency of GC alleles was above 0.5 in a majority of species, which is not expected from sequences at mutational equilibrium (Glémin et al. 2015). We applied the exact same analysis to UTR sequences and found that the mean allele frequency of GC alleles in UTRs was strongly correlated to the mean allele frequency of GC alleles at synonymous sites across species (figure 1a). This shows that the evolutionary fate of AT→GC and GC→AT mutations at third codon positions is primarily governed by forces similarly impacting non-coding DNA, i.e., independent of selection on codon usage. We reproduced the analysis using G↔C and A↔T polymorphisms instead of AT↔GC ones and obtained a very different picture (figure 1B): the frequency of C and T alleles were close to 0.5 in all species and showed no significant correlation between UTRs and third codon positions. Then we correlated GC3 and GC_UTR to gene expression level and obtained significant relationships, either positive or negative, in a majority of species (Table S1). The hypothesis of selection on codon usage bias does not predict any relationship between GC_UTR and expression. These results rather suggest that our data are probably affected by the well documented GC-bias in Illumina libraries (Benjamini and Speed 2012).

**Figure 1.**
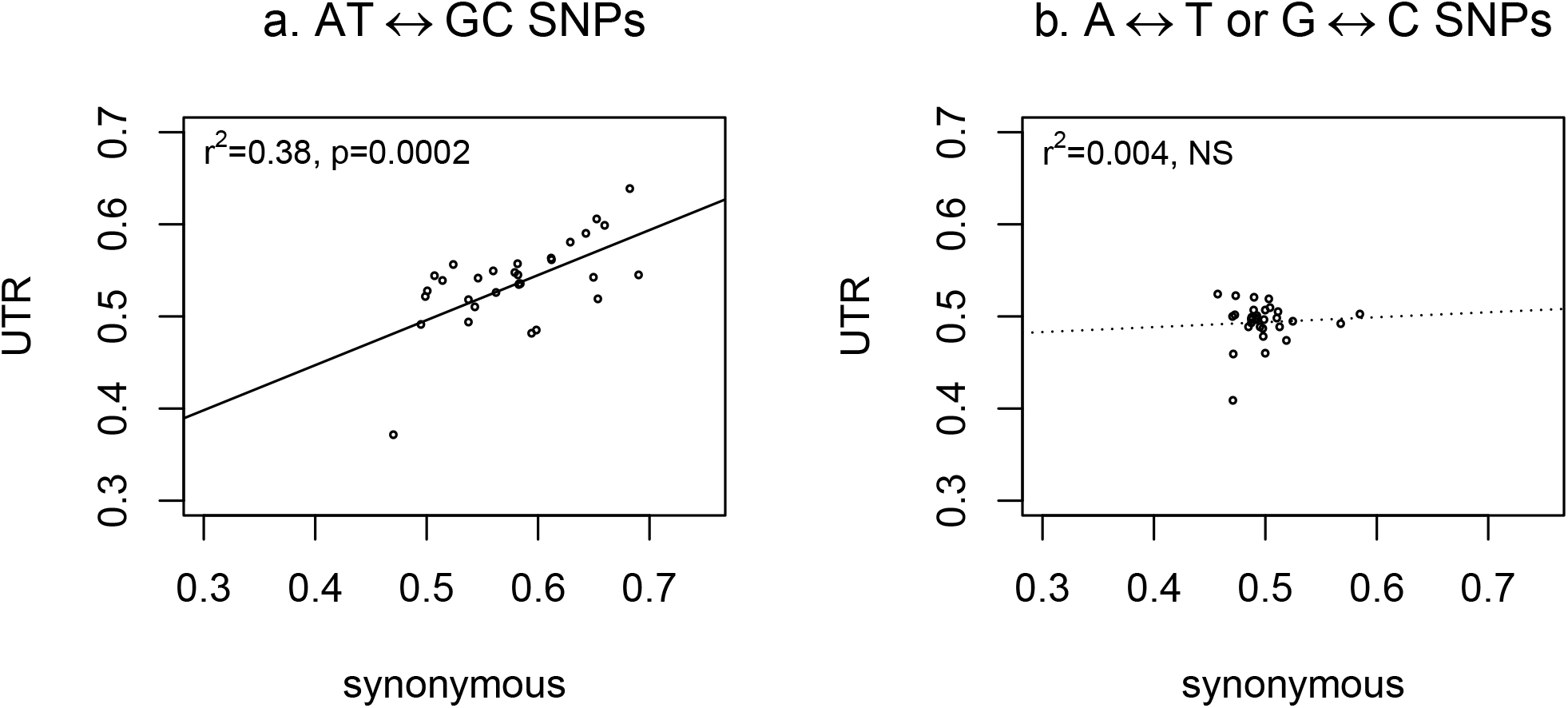
Mean allele frequency at GC-changing and GC-conservative SNPs. Each dot represents a species. X-axis: synonymous SNPs; Y-axis: flanking non-coding SNPs. a. Mean allele frequency of G or C alleles at AT vs. GC SNPs. b. Mean allele frequency of T or C alleles at A vs. T and G vs. C SNPs.

We concluded from this analysis that (i) GC-content evolution in animals involves complex processes and departs the mutational equilibrium, in agreement with the gBGC hypothesis, and (ii) the dynamics of AT↔GC mutations and the relationship between GC-content and sequencing depth are likely to confound inferences of selection on codon usage bias. To solve this problem, we decided to separately analyse two categories of mutations. Synonymous codon pairs of the form XYA / XYT or XYG / XYC were used to analyse selection on codon usage bias. There are 17 such GC-conservative pairs of synonymous codons in the standard genetic code – two per four-fold or six-fold degenerate amino-acid, and the isoleucine-coding ATA / ATT pair. The other pairs of synonymous codons, which differ by an AT↔GC change, were separately analysed to characterize the impact of GC-content dynamics and gBGC on the genomes of animals.

### Translational selection on codon usage

For each pair of GC-conservative codon and each CDS, we calculated the relative frequency of the C- or T-ending codon as

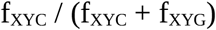

or

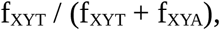

where f_XYZ_ is the frequency of occurrence of codon XYZ in the considered CDS (X, Y and Z in {A;C;G;T}). We correlated these with gene expression level. The corresponding correlation coefficient, hereafter called *r_PYR_* (for pyrimidine), were taken as measures of preference for codons XYC or XYT, compared to XYG or XYA. This was done independently in 17 GC-conservative codon pairs and 31 species, i.e., 527 correlation analyses. 454 of the 527 estimated *r_PYR_* (86%) were positive, indicating a general preference for C over G and for T over A at synonymous positions in animals. *r_PYR_* was above 0.1 in 50 codon pair analyses from 11 distinct species. In only two analyses did the correlation coefficient take values below −0.1, indicating a clear preference for A – codons CCT/CCA (Pro) and GGT/GGA (Gly) in nematode *Caenorhabditis brenneri*. No value below −0.1 was found for XYC vs XYG codon pairs.

We analysed polymorphism data in order to test whether differential codon usage between highly expressed and lowly expressed genes reflects the action of natural selection. For each pair of GC-conservative, synonymous codons, we extracted the corresponding biallelic SNPs and recorded *MAJ*, the binary variable equal to one when the frequency of the C or T allele was above 0.5 and zero when it was below 0.5, SNPs at which allele frequency was exactly 0.5 being disregarded. This was done independently in the 31 species. Then we pooled the data across codons and species and performed a logistic regression of *MAJ* on the correlation between codon usage and expression, *r_PYR_*.

We found a highly significant, positive effect of *r_PYR_* on *MAJ* (p-val<10^-15^), indicating that synonymous codons more commonly used in high-expressed genes tend to segregate at high population frequency, consistent with the hypothesis of translational selection on codon usage. The data set was split in four bins of species defined on the basis of π_N_/π_S_, a variable negatively correlated to *N_e_* (Romiguier et al. 2014), and the logistic regression was separately applied to the four bins. The relationship was strongly significant as far as the two low-π_N_/π_S_ bins were concerned (p-val<10^-12^), less strongly so in the third bin (p-val<10^-8^), and not significant in the high- π_N_/π_S_ bin. Translational selection on codon usage, therefore, is apparently affected by variations in *N_e_* among species. When species were analysed separately, a significant relationship between allele frequency and *r_PYR_* was detected in twelve species (Table S1). The mean π_N_/π_S_ of these twelve species was 0.089, while the mean π_N_/π_S_ of the 19 species for which no significant effect was detected was 0.15. The two groups of species also markedly differed in terms of average propagule_size (0.69 vs. 8.1 mm) and average longevity (7.0 vs 34 years)

The *N_e_* effect on codon usage bias is illustrated in figure 2a, in which we plotted the average allele frequency of XYC or XYT codons against *r_PYR_*, separately in low π_N_/π_S_ (top) and high π_N_/π_S_ (bottom) species. Codon pairs for which less than 20 SNPs were available in the considered species were here excluded. Figure 2a shows a strong, positive correlation between codon usage bias and allele frequencies in low-π_N_/π_S_, large-*N_e_* species. Red dots in figure 2a correspond to the nematode *C. brenneri*, in which codon usage bias is particularly pronounced. *C. brenneri* has the lowest π_N_/π_S_ of the whole dataset. In contrast, no such relationship was uncovered in high-π_N_/π_S_, small-*N_e_* species (figure 2b).

**Figure 2.**
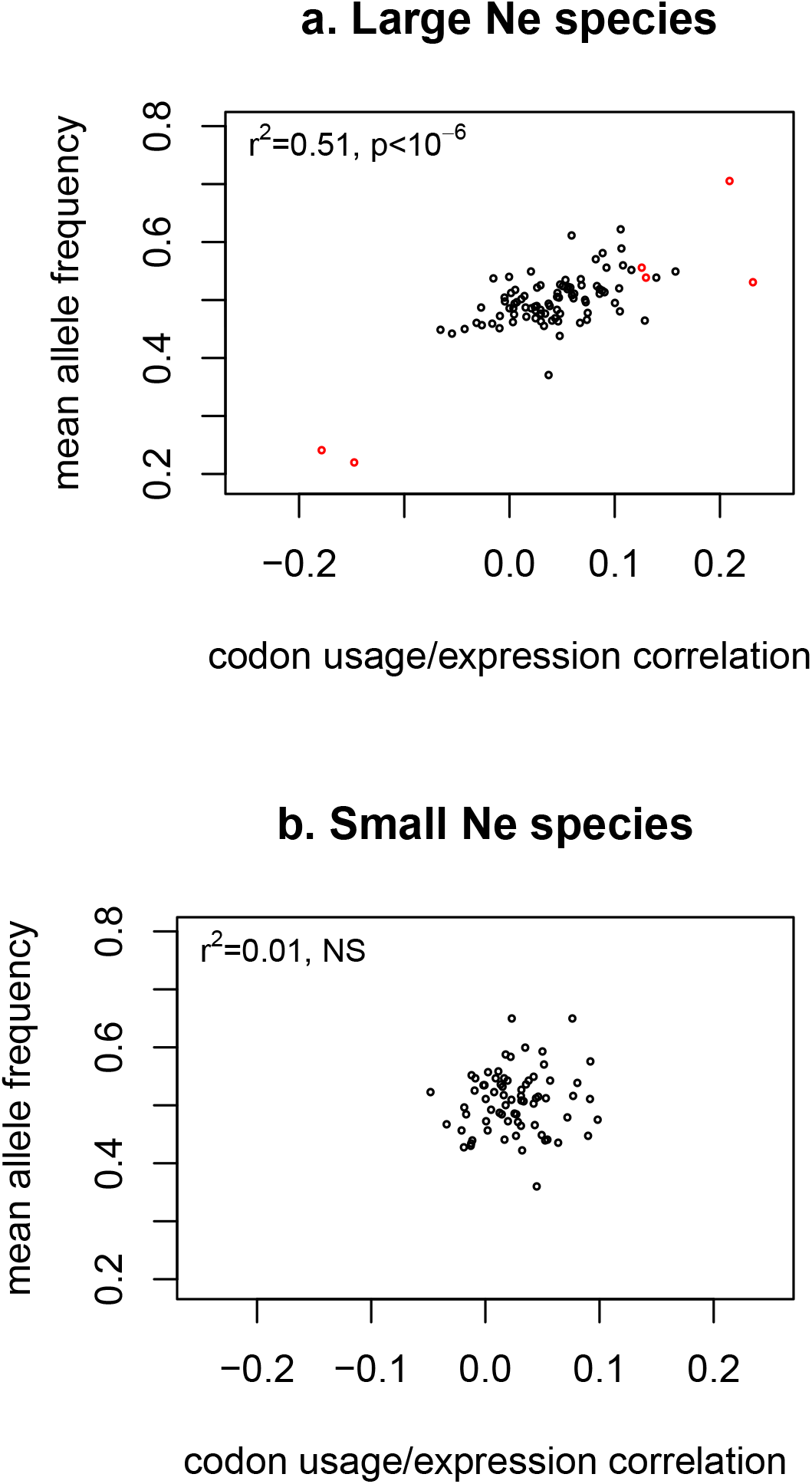
Intensity of selection on codon usage bias. Each dot represents a particular pair of synonymous codon in a particular species. Only GC-conservative pairs of synonymous codons for which a minimum of 20 SNPS are available are considered. X-axis: *r_PYR_*, the correlation coefficient between C or T usage and gene expression. Y-axis: mean frequency of the C or T allele. a. Species in which π_N_/π_S_<0.08. b. Species in which π_N_/π_S_>0.12. Red dots correspond to the nematode *Caenorhabditis brenneri*.

Not the same tissue has been used for RNA extraction in distinct species of our sample, and this might affect our results. In particular, our estimate of gene expression level could be less relevant in species where RNA has been extracted out of a single tissue, compared to multiple tissues. To control for this problem we focused on the subset of 20 species in which RNA had been extracted from at least four distinct tissues or the whole animal body (Table S1). We reproduced the above analyses and obtained very similar results (figure S1), indicating that our report of a link between π_N_/π_S_, codon usage and gene expression is not affected by tissue choice.

It has been suggested that growth rate, not *N_e_*, could be the main driver of codon usage bias intensity across species (Subramanian et al. 2008). The two effects are not easy to disentangle since *N_e_* was found to be strongly correlated to life history traits related to growth rate, such as fecundity, longevity and propagule size, in animals (Romiguier et al. 2014). To address this problem we focused on insects and compared eusocial with solitary species. Eusocial species are characterized by a dramatic reduction in *N_e_*, compared to solitary insects (Romiguier et al. 2014b). Eusocial and solitary insects, however, share similar cellular and developmental processes, so that the selective pressure for efficient protein translation can be assumed to be similar in the two groups of species.

We downloaded transcriptome data from 20 eusocial and 30 solitary insects and calculated in each species the 17 *r_PYR_*, our measure of preference for codons XYC (respectively, XYT). 691 (81%) of the 850 estimated correlation coefficients were positive, similarly to our main data set. We considered the 850 *r_PYR_* as independent data points and tested the effect of eusociality on the absolute value of this variable (figure 3a). We found that translational selection on codon usage is significantly stronger in solitary than in eusocial insects (t-test, p-val<10^-15^). Eusocial insects belong either to Hymenoptera (ants, eusocial bees and eusocial wasps) or to Blattodea (termites). To control for taxonomy the data set was split in five categories: eusocial Hymenoptera, solitary Hymenoptera, eusocial Blattodea, solitary Blattodea, and other solitary insects (figure 3b). The effect of eusociality on *r_PYR_* was significant both within Hymenoptera (t-test, p-val<10^-3^) and within Blattodea (t-test, p-val<10^-3^).

**Figure 3.**
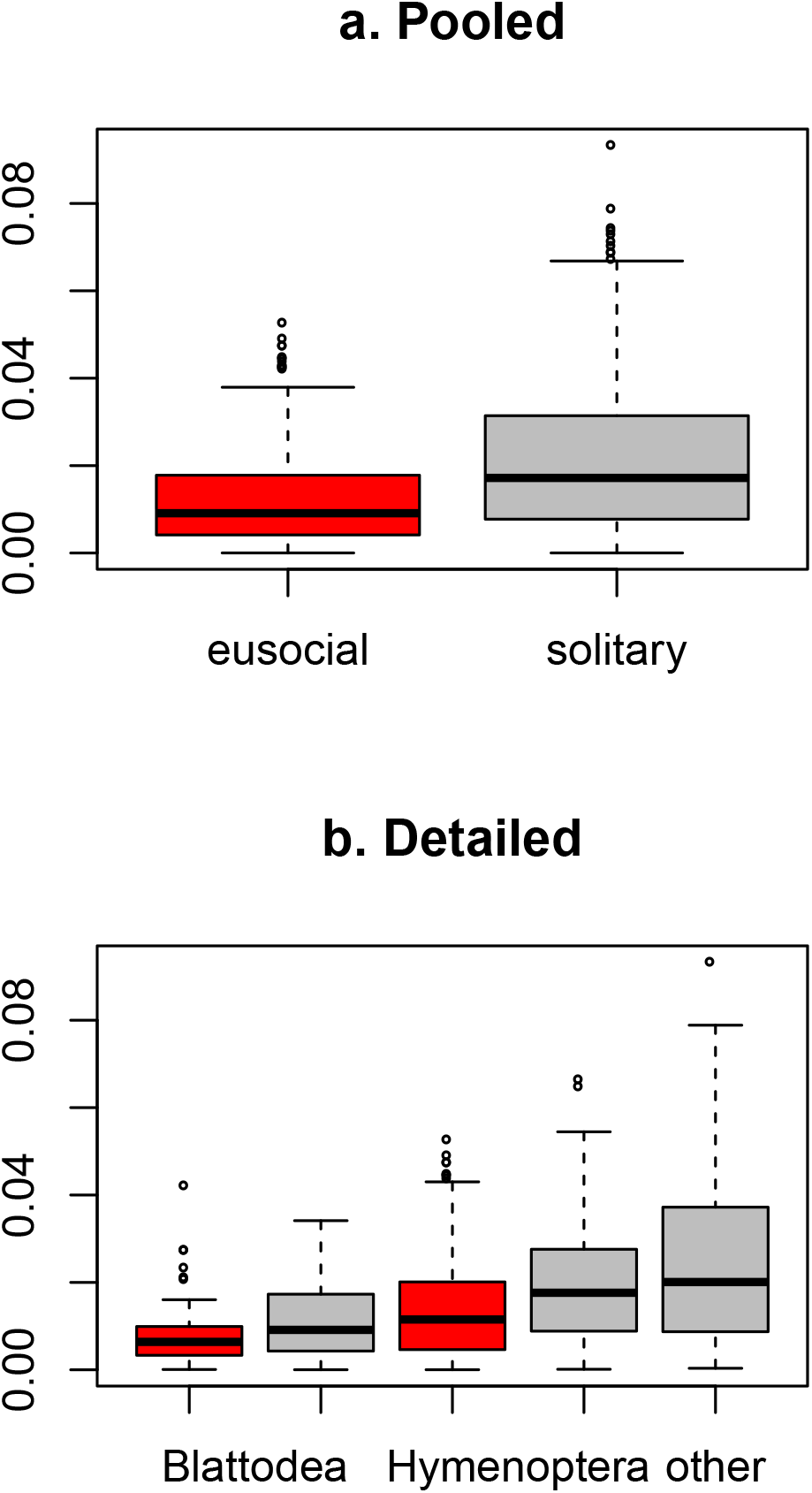
Codon usage bias in eusocial vs. solitary insects. a. Distribution of the absolute value of *r_PYR_*, the correlation coefficient between C- or T-ending codon frequency and gene expression, in 20 species of eusocial insects (red) vs. 30 species of solitary insects (grey). b. Eusocial species (red) are split in Hymenoptera vs. Blattodea; solitary species (grey) are split in Hymenoptera vs. Blattodea vs. other.

### GC-biased gene conversion analysis

In each species synonymous SNPs were oriented by assuming that the outgroup species carries the ancestral allele. We focused on the 28 species for which at least 500 oriented synonymous SNPs were available. We calculated the frequency of the derived alleles and found that in 27 species out of 28 the average frequency of alleles resulting from an AT→GC mutation was above the average frequency of alleles resulting from a GC→AT mutation (Table S1), in agreement with the gBGC hypothesis. The unfolded SFS for AT→GC SNPs, GC→AT SNPs, and GC-conservative SNPs, respectively, were built. A mutation/drift/gBGC model was fitted to the three SFS (Glémin et al. 2015). A significantly positive segregation bias in favour of GC alleles was detected in 17 species out of 28 (Table S1, figure 4a). These species belong to six of the eight metazoan phyla that were sampled. In one species, *Eunicella cavolinii* (gorgonian), the model returned significant support for a segregation bias in favour of A and T. The estimated scaled gBGC coefficient, *B*, was not significantly correlated with π_N_/π_S_, propagule size, or longevity across species. SNP sample size was limiting in this analysis, as illustrated by the large among-species variance of estimated *B* when the number of SNPs was below 5000 (figure 4a).

**Figure 4.**
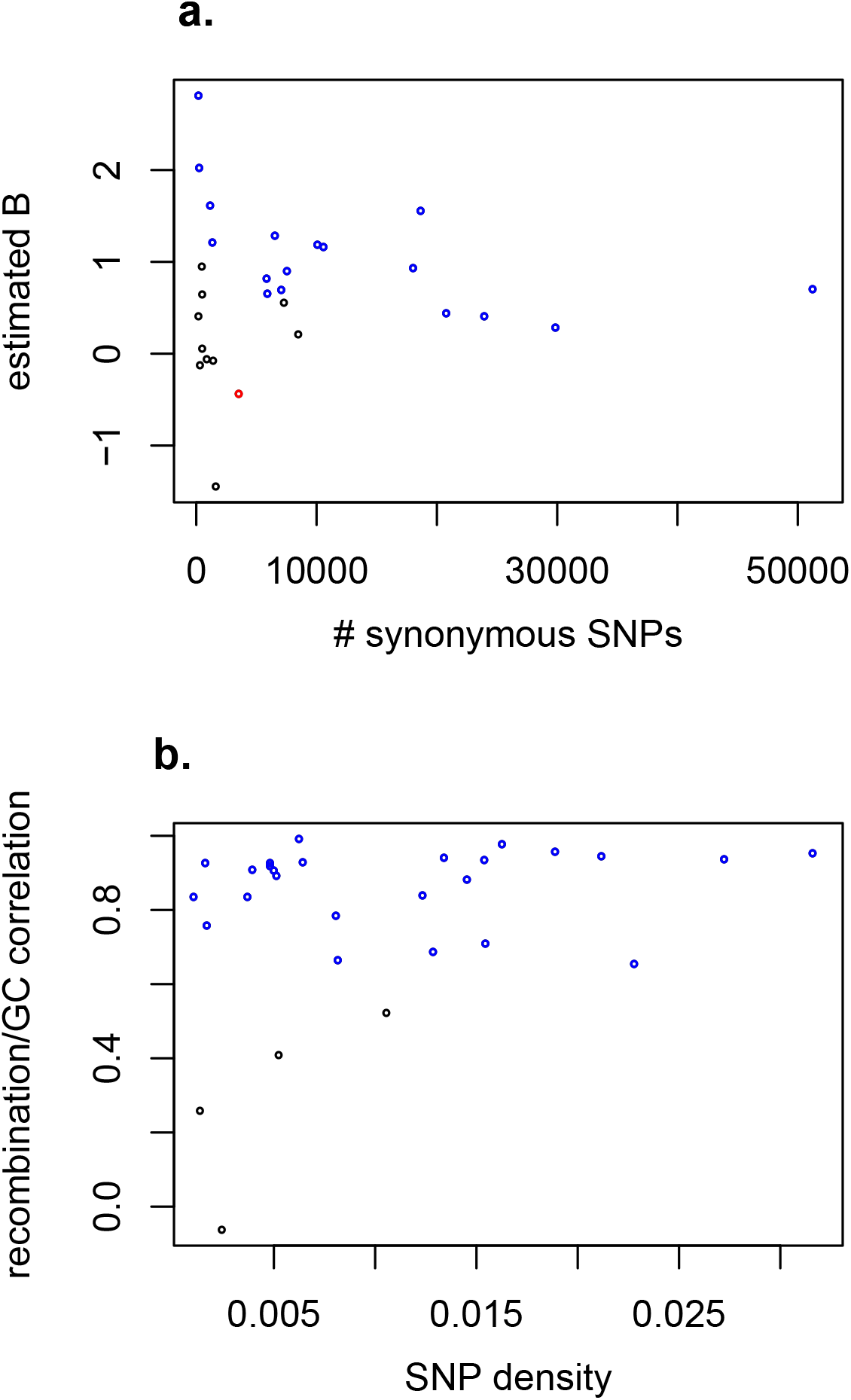
Evidence for GC-biased gene conversion. Each dot represents a species. Top: SFS analysis; X-axis: number of oriented synonymous SNPs. Y-axis: estimated scaled gBGC coefficient *B*; blue: significant, positive *B*; red: significant, negative *B*; grey: *B* not significantly different from zero. Bottom: GC/recombination relationship; X-axis: average SNP density; Y-axis: correlation coefficient between bins of *R_fgr_* and bins of GC; blue: significant, positive correlation coefficient; grey: correlation coefficient not significantly different from zero.

We investigated whether the GC-bias likely reflects the action of gBGC by analysing the impact of recombination rate. We binned contigs according to their GC-content, and measured the average recombination rate (*R*_fgr_) within each bin. We then correlated average *R*_fgr_ to average GC-content across bins in 29 species for which at least 800 loci were eligible for recombinant haplotype detection. The correlation coefficient was positive in 28 species, above 0.75 in 21 species and significantly positive (p-val<0.05) in 25 species from eight distinct phyla (Table S1, figure 4b). In one species (harvester ant *M. capitatus*) the correlation coefficient was negative but not significantly different from zero. The correlation coefficient between *R*_fgr_ and GC-content was not significantly correlated with π_N_/π_S_, propagule size, or longevity across species.

## Discussion

### Selection vs. gBGC: methodological aspects

Disentangling the effect of natural selection from that of neutral forces such as gBGC is a difficult task (Ratnakumar et al. 2010). Clément et al. (2017) introduced a model of codon usage accounting both for gBGC, which is assumed to affect AT↔GC mutations, and selection, which is assumed to affect mutations between preferred and non preferred codons. Their approach was successful in discriminating between the two evolutionary forces in eleven species of plants (Clément et al. 2017). Here we faced the additional problem that in a substantial number of species gene GC content was correlated to gene expression irrespective of codon usage, so that even defining preferred codons was challenging. We therefore addressed the problem by restricting our analysis of selection on codon usage to GC-conservative pairs of synonymous codons, which are supposedly unaffected by gBGC. One consequence is that we did not analyse two-fold degenerate codons, and among four-fold and six-fold did not compare all synonymous codons with each other, thus potentially missing a part of the signal. Of note, the classical approaches, which focus on the comparison between preferred and non-preferred codons, also have their limitations in that, as far as four-fold and six-fold codons are concerned, the non-preferred category is a mixture of several codons between which no distinction is made. For instance, published analyses of codon usage bias in *Drosophila* have hardly considered the preferences between T-ending over A-ending codons at four-fold sites, since in this group all such codons fall in the non-preferred category (but see Zeng 2010).

### Selection on codon usage in animals: a global picture

We report a significant effect of translational selection on codon usage in animals. Codons showing a higher prevalence in highly expressed genes tend to segregate at higher population frequency. We found that, with few exceptions (codons CCA and GGA in *C. brenneri*), C-ending codons are preferred over G-ending codons, and T-ending codons are preferred over A-ending. This is, to our knowledge, the first report of a general preference of pyrimidines over purines at third codon positions in animals. We checked from previously published data (Duret and Mouchiroud 1999, Lynch et al. 2017) that the trend is also found in *C. elegans* (with the same exceptions as in *C. brenneri*), *D. melanogaster*, and *D. pulex*. We would expect codon preference to be quite stable over evolutionary times since switching to a new preferred codon should impose a high genetic load by simultaneously modifying the selection coefficient at many synonymous positions. This does not explain why pyrimidines would be preferred over purines, though. In *C. elegans* and perhaps more generally, C-ending and T-ending synonymous codons are translated by the same tRNA – the so-called wobble effect – whereas each A-ending and G-ending codon has its specific tRNA (Duret 2000, Percudani 2001). For this reason one should probably expect correlated preferences for C and T at third codon positions – i.e., a frequent usage of both C- and T-ending codons when their shared tRNA is abundant, infrequent usage otherwise. Again, we see no obvious reason why the existence of a shared tRNA for C- and T-ending codons would explain that these are generally favoured over A- and G-ending codons.

The effect of translational selection on GC-conservative codons is significant but weak, and only detectable in a subset of species. This is perhaps surprising knowing that strong phenotypic effects of codon usage on expression levels of single genes have been experimentally reported in various systems, including fruit flies (Carlini et al. 2001, Carlini and Stephan 2003). Our approach, however, relies on polymorphic sites and can only detect relatively weak effects – sufficiently weak such that deleterious alleles are segregating in natural populations. Our results are indeed consistent with the existence of a broad distribution of fitness effect of synonymous mutations. In *C. brenneri*, for instance, no biased usage or skewed allele frequency distribution was detected for the proline-coding CCC vs. CCG pair, whereas strong effects were detected for, e.g., GCT vs. GCA (Ala) and CGT vs. CGA (Arg). Besides such differences between synonymous codon pairs, the effect of a particular type of synonymous mutation should also vary depending on which gene and which position is affected (Machado et al. 2017).

### gBGC is widespread across the metazoan phylogeny

We detected a significant effect of gBGC in a majority of species of the data set. gBGC manifested itself via a higher average allele frequency of GC over AT alleles both at synonymous and flanking regions, a significant difference between AT→GC and GC→AT oriented SFSs, and a correlation between the long term recombination rate and GC-content. Here scaled recombination rate was approached at contig level using an approximate method derived from the four-gamete rule. The approach is suboptimal in several respects. First, we analyse spliced sequences and have no information on intron length, so that normalization by contig length is inexact. Secondly, we analysed unphased data, thus loosing power compared to data sets consisting in experimentally phased haplotypes. Thirdly, the calculation only partially accounts for allele frequencies and the probability of detecting recombinant haplotypes when they exist – e.g., *R*_fgr_ can only increase as sample size increases. Despite these many approximations, a strong and significant correlation between *R*_fgr_ and GC-content was identified in >80% of the species we sampled, which is indicative of a prominent and widespread effect of gBGC in animals. SFS analysis corroborated this finding in uncovering a significant segregation bias in favour of G and C alleles in a majority of species. Among the 29 species for which sufficient polymorphism data was available, 28 yielded evidence for gBGC in either the SFS or the recombination rate analysis – only in the oyster Ostrea *edulis* did both approaches fail to identify a significant signal. Of note, our SFS analysis captures the effect of both gBGC and, potentially, selection on GC-ending vs AT-ending codon usage. UTR sequence analysis demonstrates the impact of gBGC (figure 1), but selection on GC-changing synonymous mutations might also be at work in some or many of the analysed species.

In animals gBGC had so far been identified in vertebrates (e.g. Figuet et al. 2014, Mugal et al. 2015), bees and ants (Kent et al. 2012, Wallberg et al. 2015), and *Daphnia* (Keith et al. 2017), but not in *D. melanogaster* (Robinson et al. 2014), albeit on the X chromosome (Galtier et al. 2006, Haddrill and Charlesworth 2008). We here considerably expand the range of species and taxa in which gBGC is documented, adding annelids, echinoderms, tunicates, nemertians, cnidarians, lepidopterans, gastropod and bivalve molluscs, decapod and isopod crustaceans. gBGC is obviously widespread among animals. It significantly impacts the population frequency and fixation probability of AT↔GC mutations in a majority of species and should be considered as a potential confounder of molecular evolutionary studies, particularly studies of molecular adaptation, not only in mammals and vertebrates (Ratnakumar et al. 2010), but more generally in Metazoa.

### Why a N_e_ effect on codon usage bias but not on gBGC?

We detected evidence for translational selection on codon usage only in the low-π_N_/π_S_, large-*N_e_* fraction of the species we sampled. The estimated efficiency of selection was strongest in the nematode *C. brenneri*, whereas no evidence for translational selection on codon usage was found in large mammals, birds, reptiles. This is in agreement with the nearly neutral theory of molecular evolution (Ohta and Gillespie 1996). Of note, the distinction between small-*N_e_* and large-*N_e_* species does not perfectly fit the vertebrates/invertebrates contrast: we detected significant evidence for translational selection on codon usage in common vole *Microtus arvalis* (π_S_=0,85%), but not in cuttlefish *Sepia officinalis* (π_S_=0,21%), for instance.

It has been suggested that among taxa variation in codon usage bias intensity is determined by variation in selective pressure, not *N_e_*, with short generation time, high growth rate species having stronger requirement for efficient protein synthesis (Subramanian et al. 2008). To distinguish between the two hypothesis we compared codon usage bias in eusocial vs. solitary insects, which differ in terms of *N_e_* but share similar developmental processes. We found a strong effect of eusociality on codon usage bias, strongly suggesting that *N_e_*, not variable selective pressure, explains the among-taxa variation we detect. Subramanian (2008) noted that under the *N*_*e*_ hypothesis one would expect a step-like relationship between codon usage intensity and *N_e_*, since the theory predicts no biased usage for every *N_e_* well below 1/4*s* and almost perfect codon usage (i.e., ~100% of preferred codons) for every *N_e_* well above 1/4*s, s* being the selection coefficient in favour of preferred codons. This rationale, however, implicitly assumes that a constant selection coefficient applies to every synonymous mutation, which is unlikely to be true. If one rather assumes a distribution of *s* across synonymous mutations then a gradual effect of *N_e_* on the intensity of codon usage bias is expected, consistent with our and Subramanian’s (2008) results.

In contrast, no relationship was detected between the intensity of gBGC and *N_e_* in our analysis. Strong, significant gBGC was detected in the high π_N_/π_S_, presumably small-*N_e_ Lepus granatensis* (hare) and *Abatus cordatus* (brooding sea urchin), for instance, while the effect was weaker in the low-π_N_/π_S_, presumably large-*N_e_ M. galloprovincialis* (mussel, Table S1). A similar pattern was recently reported in plants, based on a data set of eleven species (Clément et al. 2017). This result is somewhat surprising in that, just like selection, gBGC should only be effective if of magnitude well above that of drift. The intensity of the signal for gBGC is expected to be determined by the product of four parameters, namely *N_e_*, the effective population size, *r*, the per base recombination rate, *l*, the length of gene conversion tracts, and *b*_0_, the repair bias in favour of GC. The *rlb*_0_ product is often denoted as *b* (e.g. Glémin et al. 2015). Our results rule out the hypothesis that *b* is constant –or a *N_e_* effect should be detected. *r, l* and/or *b*_0_ must therefore vary substantially across species, and/or be inversely related to *N_e_*.

The average per base recombination rate is known to vary among taxa (e.g. Wilfert et al. 2007). Based on our data, however, we recovered a strong, positive relationship between π_S_ and average *R_fgr_* across species (r^2^=0.78, *p*-val=4.3 10^-11^). π_S_ is an estimate of 4*N_e_u*, where *u* is the per base mutation rate, and *R_fgr_* is an estimate of 4*N_e_r*. The fact that these two variables are so strongly correlated does not suggest that *r* varies sufficiently among species to erase the *N_e_* effect. There is evidence that the length of gene conversion tracts (*l*) varies across species. For instance, in mammals, *l* is of the order of 400 bp for crossovers and 50 bp for non-crossover recombination events (Cole et al. 2014), whereas *l* is about 2000 bp for both type of events in budding yeast (Mancera et al. 2008). In drosophila, non-crossover gene conversion tracts are on average 440 bp long (Miller et al. 2016), i.e. about 8 times longer than in mammals. More data are crucially needed to characterize more thoroughly the variation in *l* among animals. Repair bias *b*_0_, finally, was estimated to be 0.014 in yeast (Mancera et al. 2008), 0.12 in *Daphnia* (Keith et al. 2016), 0.18 in flycatcher (Smeds et al. 2016), and up to 0.36 in humans (Halldorsson et al. 2016). This so far limited sample suggests that *b*_0_ varies substantially among species and could be inversely correlated with *N_e_*, perhaps explaining the absence of a *N_e_* effect on gBGC intensity in our analysis.

We can think of two possible reasons why *b*_0_ would scale inversely with *N_e_*. First, gBGC is generally speaking a deleterious process in that it promotes G and C alleles irrespective of their effect on fitness (Galtier et al. 2009, Glémin 2010, Necsulea et al. 2011). It might be that the molecular machinery involved in recombination is more efficiently selected to minimize *b* in large *N_e_* species. A formal model would be required to validate this verbal hypothesis, though. Secondly, Lesecque et al. (2013) demonstrated that in yeast, when several SNPs are part of the same conversion tract, these are most often converted in the same direction – same donor and same recipient chromosomes – the direction only being influenced by SNPs located at the extremities of tracts. This implies a mechanical decay of the average GC bias as the number of SNPs per tract increases, since AT vs. GC SNPs located in the middle of a conversion tract are converted in either direction with probability 0.5. This mechanism, if effective in animals too, might contribute to explaining the lack of a *N_e_* effect on gBGC intensity, SNP density being positively correlated with *N_e_*.

## Conclusions

Translational selection is a significant determinant of codon usage patterns in large-*N_e_* species of animals, but is weak or absent in small-*N_e_* ones, such as large vertebrates and social insects. In contrast, gBGC is widespread across animals and of strength independent of *N_e_*. gBGC is therefore a major confounder that must be seriously taken into account in any analysis of codon usage bias. This study uncovered two unexpected results that remain to be elucidated, i.e., a general preference for C- and T-ending codons over G- and A-ending ones, respectively, and an inverse relationship between the recombination-associated GC repair bias and *N_e_*.

## Acknowledgments

We thank Thomas Bataillon for helpful suggestions regarding statistics and the Montpellier Bioinformatics and Biodiversity platform for computational resources. This work was supported by European Research Council grant 232971, Swiss National Foundation grant CRSII3_160723, and Agence Nationale de la Recherche grant ANR-15-CE12-0010.

## Supplementary material

**Table S1. Species sample, major variables and results of the study**.

Header: r_VAR1_VAR2 denotes Pearson’s coefficient correlation between VAR1 and VAR2; sig_VAR1_VAR2 is the associated significance level; *: p-val<0.05; **: p-val<10^-6^; “logistic” means that logistic, not linear, regression was performed.

**Table S2. Insect species samples and IDs**.

**Figure S1. Intensity of selection on codon usage bias, multi-tissue species only** Legend: see figure 2.

